# Maternal regulation of the vertebrate egg-to-embryo transition

**DOI:** 10.1101/2023.06.22.546181

**Authors:** Ricardo Fuentes, Florence L. Marlow, Elliott W. Abrams, Hong Zhang, Manami Kobayashi, Tripti Gupta, Lee D. Kapp, Zachary DiNardo, Ronald Heller, Ruth Cisternas, Felipe Montecinos-Franjola, William Vought, Mary C. Mullins

## Abstract

Egg activation and early embryonic development are complex and highly regulated processes that involve a series of coordinated cellular and molecular events after fertilization. While significant progress has been made in unraveling the mechanisms underlying these processes, there remains a need to comprehensively understand the precise molecular pathways and regulatory factors involved. We characterized four recessive maternal-effect mutants identified from a zebrafish forward genetic screen that function during the egg-to-embryo transition. We found that these genes encompass distinct aspects of egg activation, including cortical granule biology, cytoplasmic segregation, and the suppression of microtubule organizing center (MTOC) assembly and ectopic aster-like microtubule formation. These genes are essential to the development of the early embryo and the establishment of the basic body plan. Notably, we discovered a novel gene that we named *krang,* which is highly conserved across metazoans. Maternal Krang was found to be associated with the function of cortical granules during egg activation. This collection of mutants represents valuable tools to understand the genetic architecture underlying phenotypic traits shaping the egg-to-embryo transition. Furthermore, these results highlight the evolutionary conservation of maternal functions in diverse species. By deepening our understanding of these findings, we will improve our knowledge of reproductive traits and potentially develop new diagnostic tools to address human reproductive disorders.

**Author Summary:** During egg activation, the egg undergoes a series of biochemical and cellular modifications that activate its dormant metabolism, preparing it to initiate embryonic development. Egg activation initiates the completion of meiosis, and the newly formed zygote undergoes a series of additional changes, including the cell cycle establishment. However, our knowledge of the molecular mechanisms controlling these processes remain limited. We identified four recessive maternal-effect mutants in zebrafish that exhibit a range of developmental alterations during egg activation and early embryogenesis. We found that these maternally acting mutant genes are associated with defects in cortical granule exocytosis, yolk-cytoplasm segregation, microtubule nucleation and dynamics that are critical for the egg-to-embryo transition. Our results suggest that the proper regulation of these processes is essential for successful egg development and embryogenesis. We characterized the Krang factor, which regulates aspects of egg activation likely by modulating the secretory pathway. Further studies on this gene may provide new insights into the molecular mechanisms underlying oocyte development and egg quality acquisition. Overall, our collection of maternal-effect mutants sheds light on the proper regulation of key molecular and cellular events for successful egg development, with important implications for reproductive medicine and assisted reproductive technologies.

## Introduction

The transition from the dormant egg to an embryo is a complex process that involves precise and evolutionary conserved phenogenetic associations (Abrams & Mullins, 2009; Fuentes et al., 2020; Horner & Wolfner, 2008; Schultz, Stein, & Svoboda, 2018). After the sperm and egg have fused, the resulting zygote undergoes several rounds of cell division to form the early embryo, which involves cell divisions without an increase in overall size. The successful completion of this transition is critical for the development of the embryo and dependent on the function of maternally-loaded factors in all animals examined (Abrams & Mullins, 2009; Fuentes et al., 2020; Lindeman & Pelegri, 2010; Pelegri, 2003). However, current knowledge of the expression and spatiotemporal distribution of maternal molecules regulating proper egg-to-embryo transition is poorly understood.

Egg activation is typically initiated by fertilization, while in zebrafish contact of the egg with a hypotonic solution induces activation. Following the activation stimulus, a wave of calcium release initiated at the sperm entry point triggers a multitude of events, including the exocytosis of secretory vesicles called cortical granules (CGs) and the resumption of meiosis II that is completed with extrusion of the second polar body (Leung, Webb, & Miller, 1998; Mei, Lee, Marlow, Miller, & Mullins, 2009; Sharma & Kinsey, 2008). In zebrafish, the calcium wave also triggers the cytoplasm to segregate to the animal pole to form the single cell blastodisc.

In metazoans, CGs are synthesized, transported and docked at the oocyte cortex during oogenesis (reviewed in (Wessel et al., 2001). The CGs contain enzymes, glycosylated components and structural proteins that remodel the vitelline envelope surface, driving elevation and hardening of the vitelline envelope (referred to as the chorion in zebrafish and zona pellucida in mammals) during egg activation (reviewed in (Wessel et al., 2001). CG exocytosis prevents polyspermy by modifying the vitelline envelope, which also hardens the vitelline envelope, providing protection to the prospective embryo in organisms of different taxa (reviewed in (Liu, 2011; Rojas et al., 2021; Wessel et al., 2001). Abnormalities in CG exocytosis and the vitelline envelope can cause a failure in fertilization, early embryonic development, and/or implantation (T. Rankin, Talbot, Lee, & Dean, 1999; T. L. Rankin et al., 2001; Zhou et al., 2014). Therefore, advances in understanding the molecular mechanisms governing CG exocytosis and vitelline envelope formation are also relevant to human assisted reproductive technology (ART) (Cappa, de Paola, Wetten, De Blas, & Michaut, 2018). In animal systems, cells divide in the absence of cell growth during early embryogenesis. Thus, the spatial and temporal reorganization of maternal factors into specific cytoplasmic domains within the egg and zygote prior to the first cell division, is critical for coordinating the many distinct activities carried out by the early embryo (Fuentes & Fernandez, 2010; Howley & Ho, 2000; Pelegri, 2003). In zebrafish zygotes, spatial reorganization and animal-directed flow of cytoplasm during cytoplasmic segregation form a prominent cytoplasmic domain and the site of the future embryonic cleavages, the blastodisc (Fuentes & Fernandez, 2010; Leung et al., 1998; Leung, Webb, & Miller, 2000; Shamipour et al., 2019).

Within the blastodisc, the zygotic nucleus, centrosome, and mitotic spindle form as the first cell cycle initiates (Abrams et al., 2012; Fuentes & Fernandez, 2010; Lindeman & Pelegri, 2012), in the transition from meiosis to mitotic cleavage divisions. The maternal factors guiding the spatiotemporal transport of morphogenetic determinants across the egg and zygote, and its coordination with critical events in the shift from meiosis to mitosis remain largely unknown. The zebrafish egg and zygote constitute an excellent model system to study this transition since they are easily manipulated due to their large size and external development. These features make the zebrafish suitable for the study of highly dynamic events during the egg-to-embryo transition, such as the establishment of the first cell cycle.

Here, we identified four novel zebrafish maternal-effect mutants that play a role during the transition from egg to embryo. Our forward genetic adult screen revealed mutants with unprecedented phenotypes and key functions of specific genes in egg activation. These genes have a direct impact on CG biology, cytoplasmic segregation, and the regulation of microtubule organizing center (MTOC) assembly and microtubule nucleating activity, controlling the early stages of embryonic development. Notably, we discovered the previously uncharacterized *krang* gene, which provides a foundation for future phenogenetic studies aimed at deciphering its novel function in the CG secretory vesicle pathway. Our repertoire of mutants constitutes a tool for further exploration and discovery of maternal gene functions in reproduction, and targets for potential therapies in human infertility.

## RESULTS

### Egg-to-embryo transition genes

We identified a second group of four zebrafish maternal-effect mutants with defects in the egg-to-embryo transition, *p30ahub* (henceforth called *krang^p30ahub^*)*, p08bdth* (hereinafter called *spotty^p08bdth^*), *p28tabj,* and *p26thbd* (henceafter called *kazukuram* (*kazu^p26thbd^*), which in the Mapuche language or Mapudungun means gray (kazu) egg (kuram)) (Fig. 1A; Table 1). We examined two processes of egg activation in these mutants, chorion expansion and cytoplasmic segregation. Immediately after wild-type eggs are laid and prior to their activation, the chorion is closely associated with the egg surface (Fig. 1A). Following egg activation, the chorion elevates due to CG exocytosis and the blastodisc forms by the initiation of cytoplasmic transport to the animal pole. We found that the *krang^p30ahub^* mutant showed a severe, highly penetrant small chorion phenotype (Fig. 1A). The three other mutants, *spotty^p08bdth^*, *p28tabj,* and *kazu^p26thbd^*, displayed abnormalities in cytoplasmic subcellular reorganization, subsequent cell cleavage and blastoderm formation, but did not compromise chorion elevation (Fig. 1A).

**Table 1.**
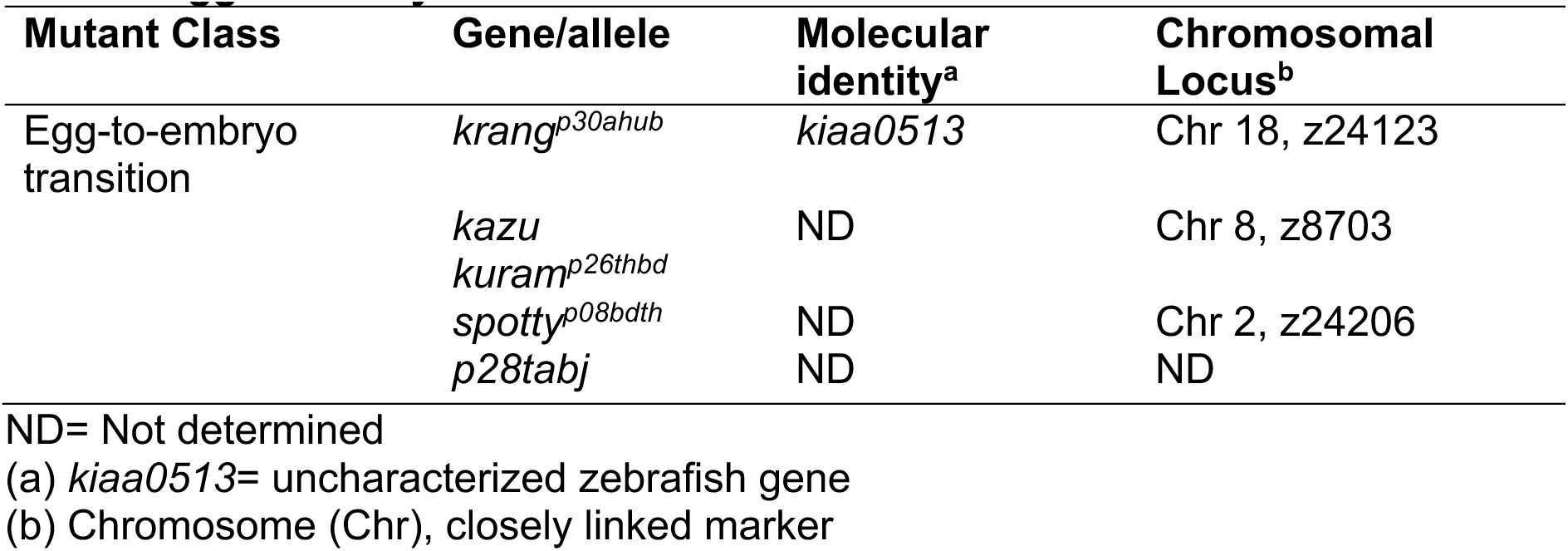
Egg-to-Embryo Transition Zebrafish Maternal-effect Genes.

**Figure 1.**
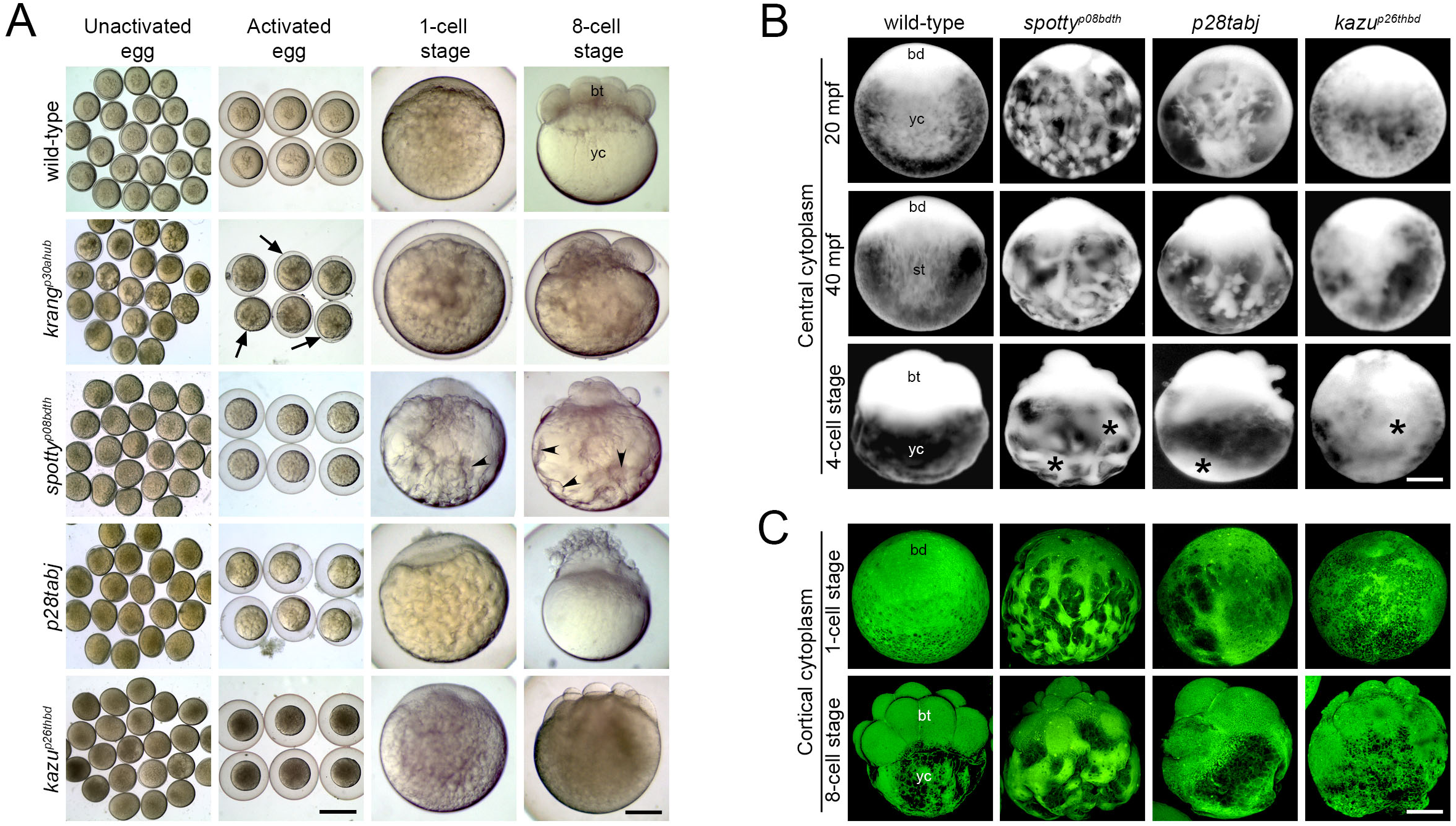
Egg activation mutants. A. Bright-field images showing early developmental stages of live wild-type and mutant eggs and embryos. All unactivated eggs collected after gently squeezing gravid females showed no detectable defects. After egg activation, however, the *krang* mutant egg displays a pronounced chorion elevation defect (black arrows). As development proceeds, the cytoplasm is abnormally distributed in the yolk cell (yc) of *spotty* mutant eggs and early embryos (black arrowheads) and a defective blastoderm (bt) is formed. *krang* (1887 eggs from 50 females)*, spotty* (1273 eggs from 40 females), and *p28tabj* (2506 eggs from 34 females) and *kazu* (1608 eggs from 32 females). **B.** Lateral view of acid-fixed wild-type and mutant eggs and early embryos showing the organization of the central cytoplasm (bright). In wild type, the blastodisc (bd) grows by animal-ward transport of cytoplasm from the yolk cell (yc) and by the 4-cell stage, the cytoplasm in the yolk cell is no longer evident. In contrast, *spotty*, *p28tabj* and *kazu* mutant eggs exhibited an abnormal distribution and severe retention of cytoplasm in the yolk cell (black asterisks). Wild-type (n=22), *spotty* (n=17), and *p28tabj* (n=19) and *kazu* (n=12). **C.** Lateral view of acid-fixed and DiOC_6_-stained wild-type and mutant eggs and early embryos showing the organization of the cortical cytoplasm (green) and yolk (dark). Wild-type (n=20), *spotty* (n=20), and *p28tabj* (n= 14) and *kazu* (n=18). Scale bar= 830 µm (A, columns 1 and 2 from left to right), 150 µm (A, columns 3 and 4 from left to right), 190 µm (B, C).

We mapped the *krang, spotty*, and *kazu* mutations to chromosomal loci and determined that *p28tabj* was not linked to the intervals of the other mutations reported here or previously (Table 1. See Material and Methods in the companion study). Based on our phenotypic and mapping analysis, we conclude that these mutations disrupt distinct genes required for chorion elevation, cytoplasmic segregation, and early embryo formation. Importantly, we identified new genes that appear to control divergent processes acting in the zebrafish egg-to-embryo transition.

### Cytoplasmic reorganization and blastoderm formation maternal egg-to-embryo mutants

In most animals with telolecithal eggs and discoidal cleavage, including zebrafish, egg activation is characterized by blastodisc elevation, which is the precursor to the blastoderm and ultimately the embryo (reviewed in (Fuentes, Mullins, & Fernandez, 2018). During blastodisc growth, the cytoplasm is transported to the animal pole through the process of cytoplasmic segregation in two main phases: the slow (20-30 minutes post fertilization, mpf) and fast (30-40 mpf) flow of cytoplasm (Fuentes & Fernandez, 2010; Leung et al., 2000). The *spotty* and *p28tabj* mutants displayed apparent normal blastodisc formation at 20 and 40 mpf, while *kazu* mutant eggs appeared normal at 20 mpf but displayed a smaller blastodisc at 40 mpf (Figs. 1A,B). For all three mutants the subsequent cell cleavage stage was morphologically abnormal, with altered cell size and organization, and the *spotty* mutant also had pockets of cytoplasm within the yolk cell (Figs. 1A,C).

To further explore the cytoplasmic segregation process, we visualized the organization of cytoplasmic domains and organelles. In acid-fixed wild-type eggs, the cytoplasm is distributed in the blastodisc and across most of the yolk cell, intermingled with YGs (Fig. 1B). As development proceeds, the blastodisc grows by transport of cytoplasm from the yolk cell via specialized transportation pathways or streamers to the animal pole (Fuentes and Fernandez, 2010). The *spotty*, *p28tabj* and *kazu* mutant zygotes showed marked changes in the organization of cytoplasm distributed in the yolk cell during the first phase of cytoplasmic segregation (Fig. 1B, 20 mpf). During the fast phase of cytoplasmic segregation when axial streamers form, the distribution of membranous organelles was also affected in the mutant zygotes (Figs. 1B, C, 40 min, 1-cell stage). Cytoplasmic patches in various regions of the yolk cell were present in these mutants but not wild-type (Figs. 1B,C). Thus, defective cytoplasmic organization and animal-ward streaming, likely explain its retention in the yolk cell in the mutant zygotes studied. This suggests that the mechanisms regulating cytoplasmic segregation are compromised in these mutants.

### Maternal Krang plays a role in cortical granule biology during egg activation

The eggs of *krang* females exhibited a fully penetrant small chorion phenotype, as determined by quantitating the extent of chorion elevation (Figs. 1A, 2A). To investigate whether the pronounced chorion elevation defect in the *krang* mutant results from a defect in CG synthesis, translocation or exocytosis, we performed MPA-staining to specifically label and track CGs. We found that CGs formed and were cortically distributed in *krang* eggs before activation (Fig. 2B). However, at 20 minutes post activation (mpa), numerous small CGs persisted in the mutant egg, along with small CG-like vesicles with reduced MPA staining within the vesicle (Figs. 2B; S1A). In addition, larger CGs were retained in the mutant subcortex, surrounded by clusters of small CGs, unlike in wild-type (Figs. 2B; S1A). By counting the number of CGs per unit area at different time points after egg activation, we found, as expected, rapid CG exocytosis in wild-type and mutant eggs (Fig. 2C). While there was a significantly reduced number of CGs in unactivated *krang* mutant eggs, the number remaining at multiple time points after egg activation was not significantly different (Fig. 2C). However, the rate of CG exocytosis was mildly retarded at 20 and 30 mpa in *krang^p30ahub^* mutants (Fig. S1B).

**Figure 2.**
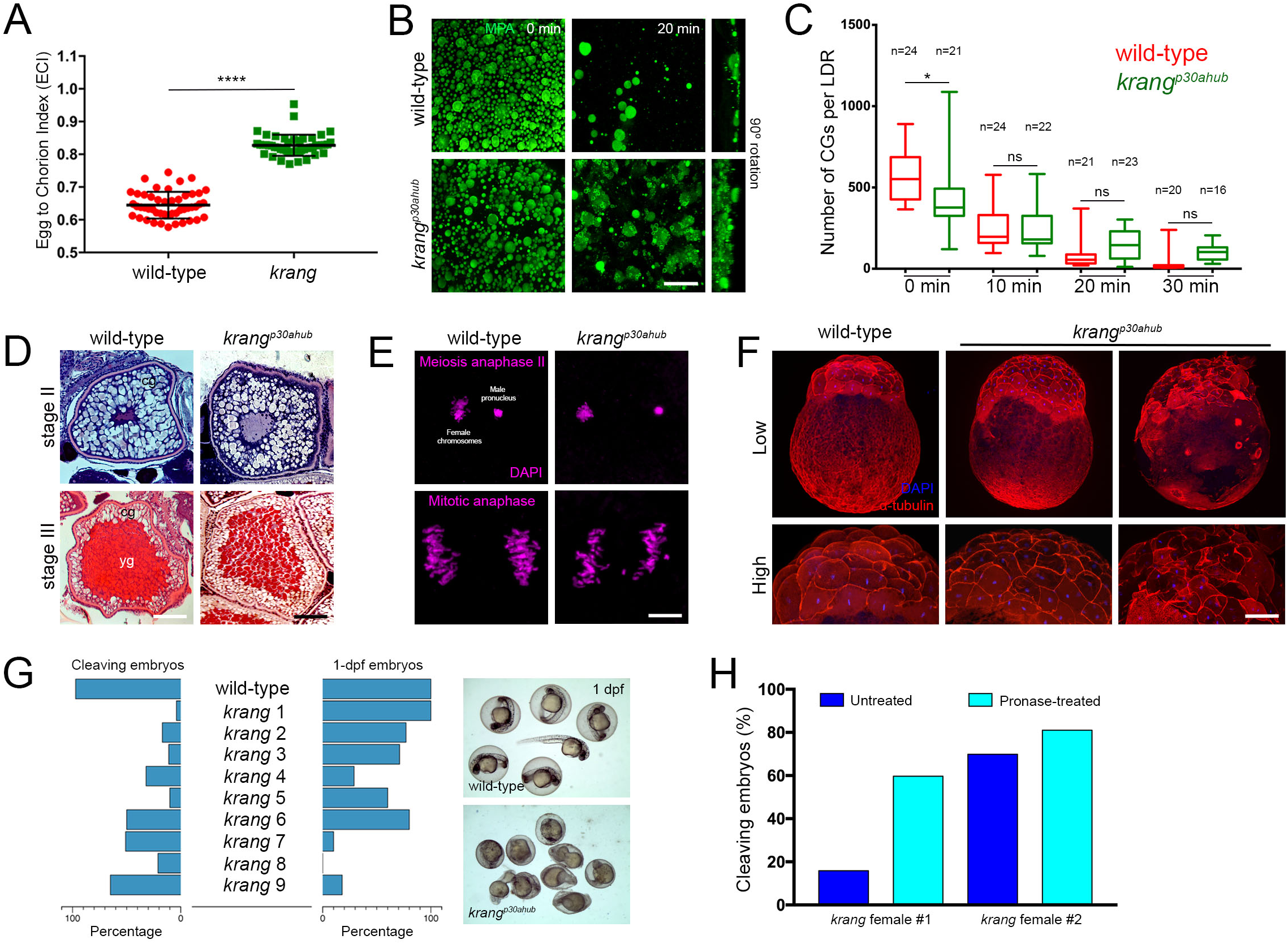
Characterization of the maternal-effect *krang* mutant phenotype. A. Quantification of the chorion elevation phenotype. Scatter plots of the measured ECI value show a penetrant chorion elevation defect in the *krang* mutant egg at 30 mpa (ECI=0.64 and 0.83; SD= 0.006 and 0.005; for the wild-type (n=48, 2 females) and mutant (n=48, 2 females) egg, respectively). Data are means ± SD. \*\*\*\**p*<0.0001 in statistical clustering analysis in a nonparametric unpaired *t test* of chorion elevation measurements. **B.** Confocal z-projections (5 μm depth) of MPA stained CGs showing their distribution in wild-type (top row, n=68 eggs from 3 females) and *krang* mutant (bottom row, n=55 eggs from 3 females) eggs at two different time points after activation. Images were taken in a lateral determined region (LDR) of the activated egg. Notice that smaller, possibly CGs lacking MPA-staining lectin accumulate in the mutant egg at 20 mpf. **C.** Box plots of CG exocytosis timing measured in the LDR of the activated egg. The average number of CGs in the *krang* mutant was significantly reduced compared to wild-type at 0 min but not significantly different at later time points. n, number of eggs used in this analysis from 3 females. **D.** Hematoxylin and Eosin stained sections of intact stage II (left column) and stage III (right column) oocytes. Wild-type (n=2 ovaries), and *krang* (n=2 ovaries) oocytes, were phenotypically comparable. CGs are formed (stage II) and accumulated at the cortex (stage III). **E.** Confocal micrographs of DAPI-stained eggs and zygotes showing normal meiosis completion and fertilization (top row, wild-type (n=14) and mutant (n=9)), and chromosome segregation during the first mitosis (bottom row, wild-type (n=13) and mutant (n=14)). **F.** Most *krang* mutant embryos failed to undergo cell divisions. Top row: wild-type embryo with a typical symmetric blastoderm at 2 hpf (left panel, n=87/93 embryos from 2 females). In contrast, *krang* mutant embryos display relatively normal (middle panel, n=78/227) or abnormal (right panel, n=147/227) blastoderm formation (n=3 mutant females). Bottom row: high magnification confocal micrographs showing the cell architecture in wild-type and mutant blastoderms. **G.** Left: Graphs showing cleavage (left) and survival (right) of cleaving wild-type and mutant embryos. Two wild-type and 9 *krang* females were analyzed. Right: Wild-type control blastula were all normal at 1 dpf (n=64/64, left panel). Most mutant embryos failed to develop beyond blastula stage (n=64/82). Mutant blastulae gave rise to 1-dpf embryos with a variable phenotype: wild-type-like (n=10/82), and reduced body axis (8/82). **H.** Bars graph showing the cleavage percentage of pronase untreated and treated *krang* embryos. Fertilized mutant eggs were subjected to pronase incubation (treated; n=125, 2 mutant females). Control cleaving percentage was obtained by incubating *krang* eggs in pronase-free medium (Untreated; n=161, 2 mutant females). SD= standard deviation; mpa, minutes post activation; dpf, days post fertilization; cg, cortical granules; yg, yolk globules. Scale bar= 40 µm (B), 25 µm (D, left column), 95 µm (right column), 50 µm (E, top row), 12 µm (E, bottom row), 160 µm (F, top row), 85 µm (F, bottom row).

To explore CG formation and localization during oogenesis, we examined ovaries from homozygous mutant females. We found that oogenesis in *krang* mutants appeared morphologically and histologically normal, where CGs were primarily formed in stage II oocytes and then translocated and docked to the cell cortex in later oogenesis stages (Fig. 2D). Collectively, these results indicate that CGs form and localize in the *krang^p30ahub^*mutant as in wild type. Possible changes in CG content or exocytosis after egg activation might impact the magnitude of chorion elevation.

We found that *krang^p30ahub^*eggs were fertilized, meiosis II resumed and completed, and the initiation of the first mitosis ensued in *krang* mutant zygotes (Fig. 2E). However, during early embryonic development, many *krang^p30ahub^* embryos failed to undergo cleavage or did so abnormally, often dying during blastula stages (Figs. 2F,G). The *krang^p30ahub^* embryos that survived to 1 dpf exhibited a wild-type phenotype, or a reduced or ventralized body axis (Fig. 2G). To test whether defective chorion elevation impairs the cleaving ability of *krang* early embryos, we evaluated the effect of chorion removal by pronase treatment after fertilization on the cleavage rate. Removal of the chorion greatly increased the fraction of cleaving embryos from mutant females (Fig. 2H).

To investigate whether defects in chorion biogenesis were associated with the chorion elevation defect, we examined its ultrastructure in cross-sectioned wild-type and *krang* whole ovaries by transmission electron microscopy. The chorion is derived from the vitelline envelope, which is made during oogenesis. In the zebrafish, the structure of the vitelline envelope consists of three zones organized horizontally around the oocyte (Hau et al., 2020; Selman, Wallace, Sarka, & Qi, 1993). In analyzing the vitelline envelope ultrastructure, we did not find any morphological variation between wild-type and *krang* oocytes (Fig. S1C). These results indicate that the chorion elevation defect, but not its biosynthesis, influences the developmental potential of mutant embryos to progress. When removed, it greatly improved cell cleavage and axis formation.

### Krang encodes a novel protein

To identify the molecular nature of the *krang* mutant gene, a combination of positional cloning and next generation sequencing strategies were carried out. We mapped the *krang^p30ahub^* mutation to chromosome 18 between markers z65576 and z10008. In testing 1380 females for recombination in the interval, we finely mapped the critical interval of the mutation to 0.87 cM between markers CR925798-1 (38.93 cM) and z58289 (39.8 cM) (Fig. 3A). We performed a genomic DNA sequence capture of the entire 1.2 Mb interval from a *krang^p30ahub^* homozygote and wild-type control (Dapprich, Ferriola, Magira, Kunkel, & Monos, 2008; Gabriel et al., 2006; Gupta et al., 2010; Nagy et al., 2007), followed by next generation sequencing of the captured DNA. Sequence analysis revealed only 3 single base pair (bp) changes within the physical interval of the mutant gene. Two of the changes were in a large first intron of a gene and the third was in the coding sequence of the same gene. This coding sequence change did not alter the Gly amino acid encoded, but was a wobble codon change from GGC to GGT. None of the changes appeared deleterious, but all were present in the same gene, *kiaa0513*. Thus, we cloned and sequenced the cDNA of *kiaa0513* from *krang^p30ahub^* and wild-type ovaries to determine if it was affected. Interestingly, sequencing of the cDNA revealed a 35 nucleotide (nt) deletion at the 3’-end of exon 3, precisely at the position of the wobble alteration (Fig. 3B). Thus, the wobble mutation (C to T) created an earlier splice donor site 35 nts upstream of the wild-type donor splice site (Figs. 3A, B), generating a transcript lacking 35 nts of the coding sequence.

**Figure 3.**
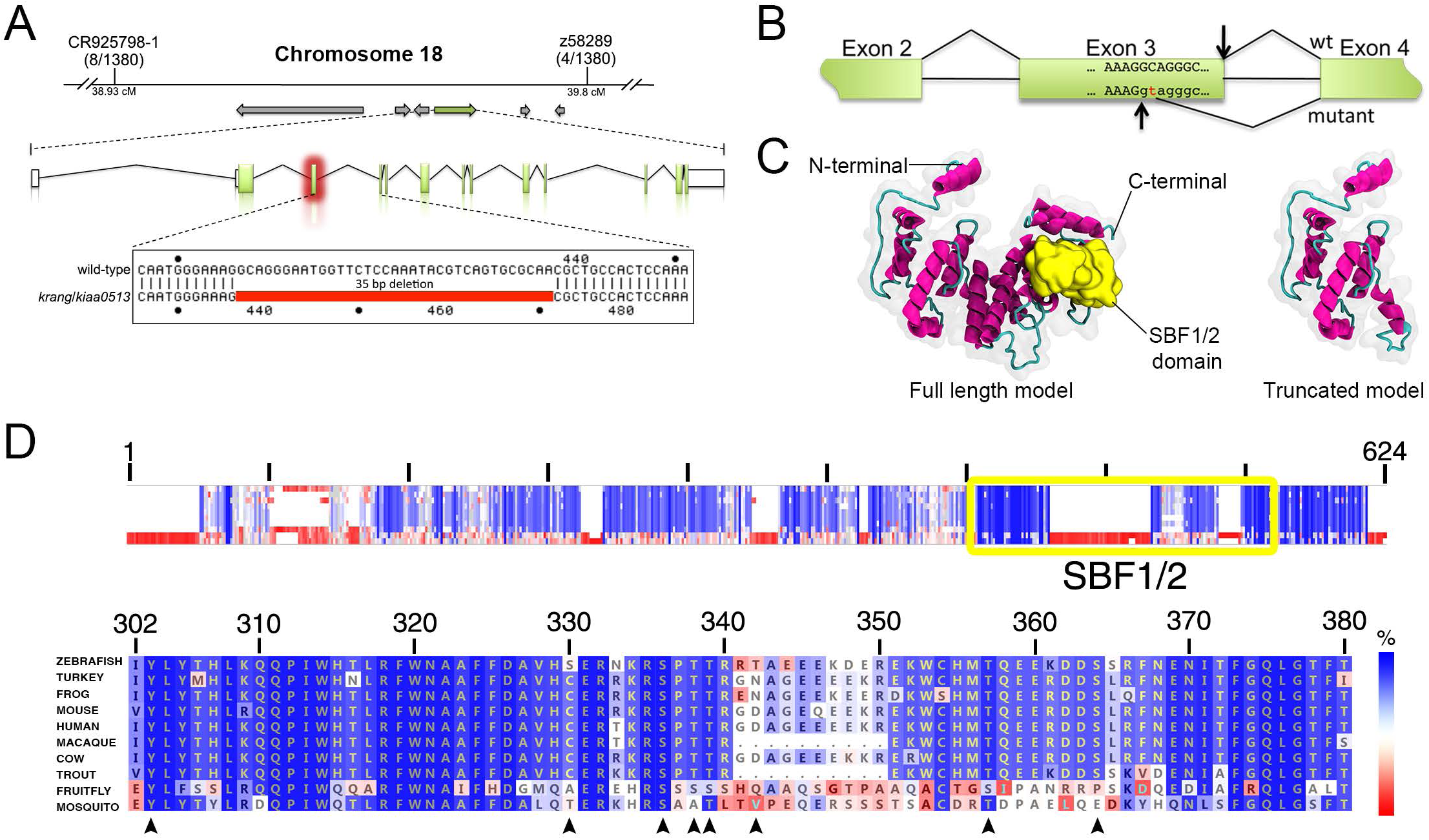
Molecular characterization of the Krang maternal factor. A. Schematic of the zebrafish *krang/kiaa0513* gene, which consists of 13 exons and 12 introns, and the 35 nt deletion in the mutant exon 3. Exons are shown as green boxes and introns as black lines. Sizes are not to scale. **B.** Schematic representation of the molecular lesion and generation of a premature donor splice site 35 nts upstream of the wild-type (wt) donor splice site in the *krang/kiaa0513* transcript. **C:** Predicted tertiary structure of the full-length zebrafish Krang protein (left) and its truncated version (right). The α-helixes and the connecting loops are colored in pink and green, respectively. N– and C-terminal domains are indicated. The SBF1/2 functional domain of the protein is shown in yellow. The approximate volumetric density map of the protein is shown in translucent gray. **D.** Krang protein sequence alignment of multiple species represented in E. Top: Schematic representation of the protein sequence alignment. The overall percentage (%) identity among sequences decreases from top to bottom. The yellow box indicates the SBF1/2 functional domain in Krang. Bottom: Detailed sequence alignment, showing high conservation of this domain in fungi, insects, fish, birds and mammals. Black arrow heads indicate conserved phosphorylation hotspots. Percent similarity is color coded by the scale bar at the right.

The *kiaa0513* transcript encodes a 424 amino acid protein with no previously known function, although highly conserved from arthropods to humans (Figs. 3C, D). The 35 nt deletion in the *krang^p30ahub^* cDNA results in a frameshift, which generates a premature stop codon following 24 aberrant amino acids (Fig. S1D). As a result, the predicted mutant protein lacks 255 of 424 amino acids, including the highly conserved SET binding factor 1/2 (SBF1/2) or myotubularin-related domain (Fig. 3C). SBF1 and SBF2 proteins (Myotubularin-related protein 5 and 13, respectively) have been previously described as homologous to phosphatases but lacking phosphatase activity and instead acting as pseudo-phosphatases, regulating the enzymatic activity of other phosphatase myotubularin-related proteins (Azzedine et al., 2003; Berger et al., 2006; Kim, Vacratsis, Firestein, Cleary, & Dixon, 2003). However, the non-catalytic phosphatase domain of these factors was not found in Krang. Amino acid alignment of zebrafish Krang to other animal orthologs revealed high similarity within the SBF1/2 domain (Fig. 3D) and phylogenetic analysis shows the conserved evolutionary relationship of Krang homologs in vertebrate and invertebrate organisms (Figs. 4A, S2). There is little information available about Krang/Kiaa0513 to date; therefore, how its SBF1/2 domain physiologically functions in egg activation will need further investigation.

**Figure 4.**
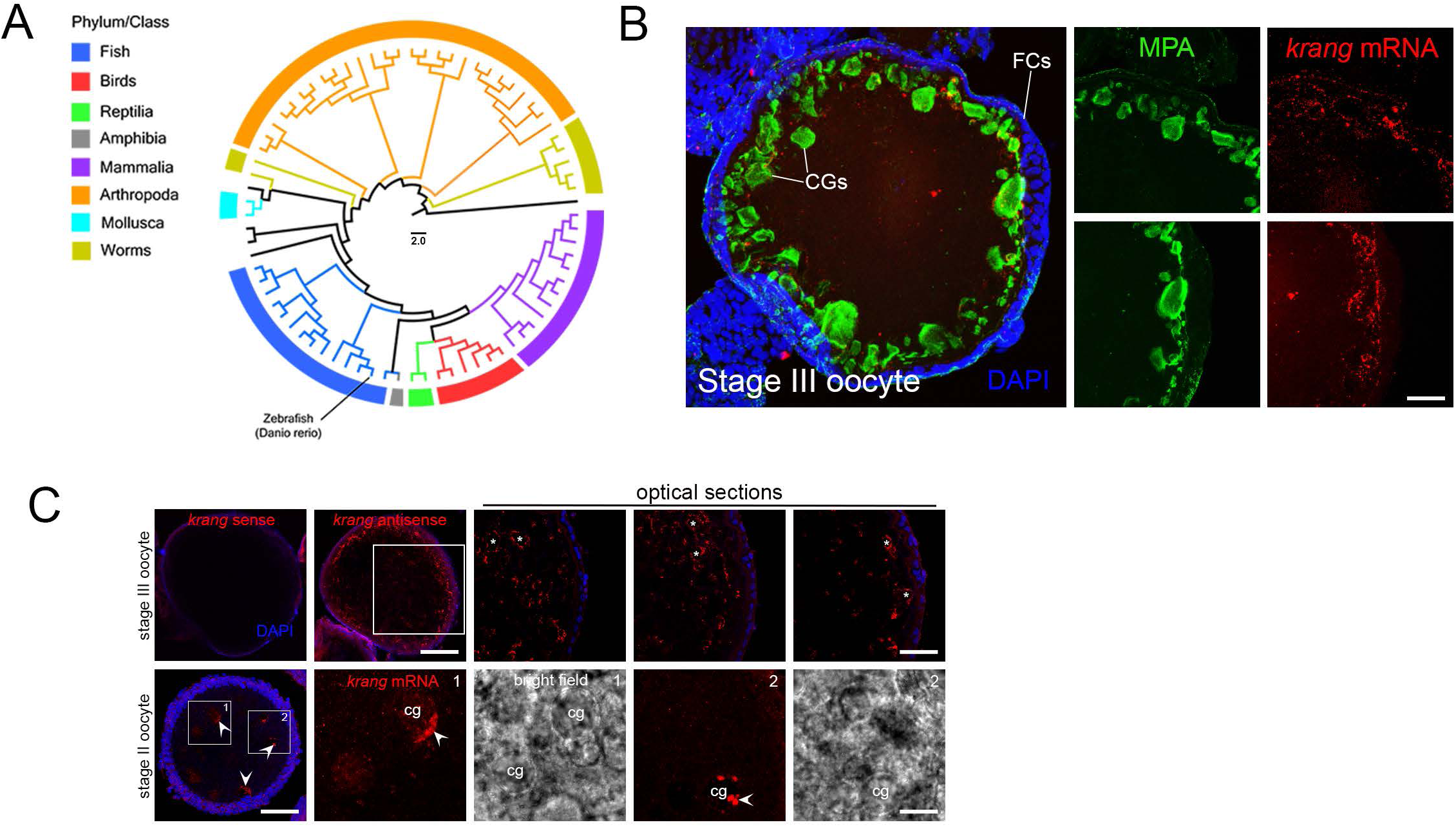
Evolutionary history and expression pattern of maternal *krang*. A. Reduced neighbor-joining phylogenetic tree of Krang homologs found in public databases indicating its evolutionary history from invertebrate to vertebrate species. The scale bar in the middle represents evolutionary distances based on residue substitutions per site. **B:** *In situ* hybridization showing the cortically-restricted distribution of *krang* mRNA in a cryosectioned stage III oocyte (n=18). Notice that the expression of the transcript is associated with MPA-stained CGs. FCs, follicle cells. **C.** Whole-mount *in situ* hybridization showing *krang* transcript localization. Top row: wild-type stage III oocyte (n=43), *krang* mRNA is peripherally distributed in the cell, presumably overlapping with CGs (white asterisks in high magnification images of the same oocyte). Bottom row: stage II oocyte (n=19), where the wild-type *krang* transcript is associated with nascent CGs (bright field and fluorescent high magnification images). Control sense probe did not show transcript signal in stage III oocytes (n=23, top row). Boxes in C show magnified areas. Scale bar= 20 µm (B), 140 µm (C, stage III oocyte), 9 µm (C, top row high magnification images), 55 µm (C, stage II oocyte), 20 µm (C, bottom row high magnification images).

To validate that *krang* is encoded by *kiaa0513*, we generated a second mutant allele, *krang^pΔ14^*, using a CRISPR/Cas9 approach. The new *krang* mutant allele is a 14 bp deletion affecting exon 5 that generates a premature stop codon (Fig. S3A). Mutant females also produced eggs wherein the chorion fails to elevate (Fig. S3B). To evaluate if *krang^30ahub^*and *krang^pΔ14^* alleles do not complement, we generated trans-heterozygous loss-of-function females (*krang^30ahub+/-^*; *krang^pΔ14+/-^*), whose eggs and early embryos displayed the *krang* chorion elevation and cell cleavage defect. In addition to the tight chorion phenotype, transheterozygous females yielded ventralized embryos, altogether showing that these two alleles do not complement and are mutations in the same gene, *krang*.

To gain insights into Krang function, we examined the expression pattern of its mRNA during oogenesis. In previtellogenic stage II oocytes, *krang* transcript puncta were primarily detected around or in nascent CGs, whereas in vitellogenic stage III oocytes, it is peripherally localized in the vicinity of CGs (Figs. 4B, C). A similar localization pattern has been described for other mRNAs encoding exocytosis and vesicular release proteins in neurons (Hobson et al., 2022). Taken together, the abnormalities observed in the mutant egg and embryo, and the spatial localization of its mRNA, suggest that Krang functions in an egg activation process, possibly regulating crucial components involved in CG dynamics and integrity, which subsequently act in chorion elevation.

### The maternal-effect kazu gene controls cell volume acquisition during embryogenesis

In cleavage stage embryos, the initial volume of cytoplasm at the one-cell stage is cleaved at each cell division in half, with no overall increase in total cytoplasmic volume until much later in development (Kane & Kimmel, 1993; Kimmel, Ballard, Kimmel, Ullmann, & Schilling, 1995). In zebrafish the extent of cytoplasmic segregation from the yolk into the blastodisc also determines the cleavage stage cell sizes. We found that the eggs of the maternal-effect *kazu^p26thbd^* mutant display a small blastodisc, and the early cell divisions were characterized by asymmetric and irregular cleavage, generating smaller and less compacted blastomeres (Fig. 5A,B). Additionally, the cytoplasm persisted within the yolk and cell detachment was also observed in *kazu^p26thbd^* cleavage stage mutants (Figs. 1B, 5A). Most *kazu^p26thbd^* mutant embryos failed to develop beyond the sphere stage and died during this period (Fig. 5A). However, those that survive generally show a reduced body axis (Fig. S4A). These results are consistent with a defect in fully segregating the cytoplasm from the yolk cell, with cytoplasm persisting within the yolk at the 4-cell stage and beyond (Fig. 1B, 5A).

**Figure 5.**
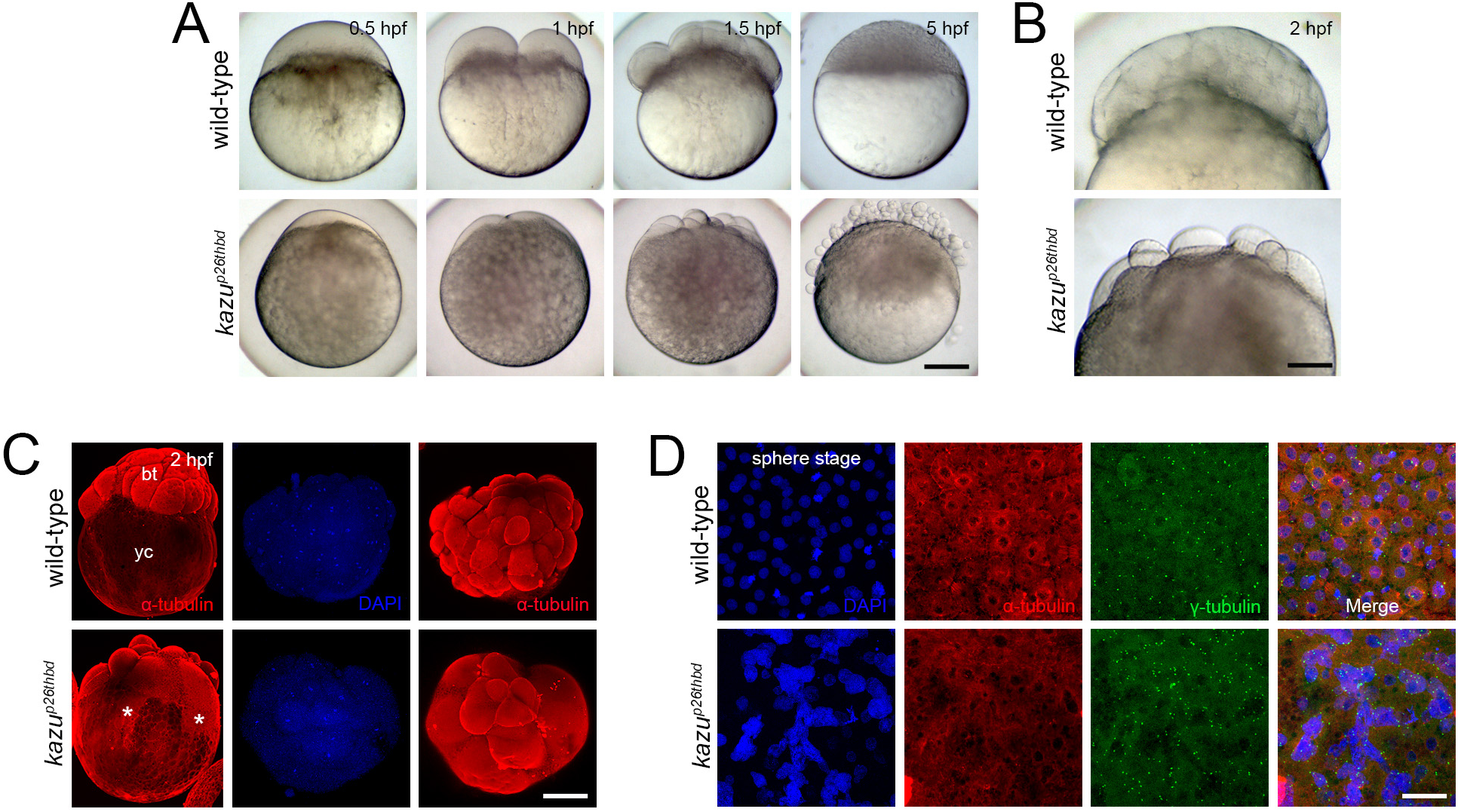
Phenotypic characterization of the *kazu^p26thbd^* mutant early embryo. A. Bright field images showing early developmental stages of whole-mount live wild-type (n=99/99 from 2 females) and *kazu^p26thbd^* (n=185/265 from 2 females) embryos. The mutant phenotype reveals a smaller blastodisc (bd) with smaller cells during cleavage. These abnormalities in cell size acquisition, together with asynchronous and unequal cell divisions, leads to the formation of a defective blastula with many detached blastomeres. **B.** Higher magnification images showing a cellularized wild-type blastoderm (n=125/125). In contrast, *kazu^p26thbd^* embryos exhibit a less compacted blastoderm containing smaller cells (n=62/91). **C.** Lateral and animal views of whole-mount wild-type (n=31/35) and mutant (n=39/47) early embryo that show the distribution of α-Tubulin at 2 hpf. In wild-type embryos, α-Tubulin is mainly distributed in the blastoderm (bt) and dividing blastomeres. In *kazu^p26thbd^* mutants, α-Tubulin is distributed in different sized cells and throughout the yolk cell (yc, asterisks). **D.** Confocal z-projections of whole-mount α– and γ-tubulin, and DAPI staining of wild-type (n=30/35) and *kazu^p26thbd^*(n=25/33) sphere stage embryos shown in animal pole views. Notice the enlargement of nuclei, multiple centrosomes and loss of microtubule structures in mutant embryos. Scale bar = 180 µm (A), 110 µm (B), 230 µm (C), 100 µm (D).

We further examined the impact of defective cytoplasmic segregation and a small blastodisc to early embryo development. Immunostaining revealed that α-tubulin distribution was affected in early mutant embryos (Fig. 5C). α-Tubulin in mutant embryos extended from the blastomeres into the yolk cell at the 16 to 32-cell stage, whereas it was restricted to the blastomeres in wild-type. In large parts of the yolk cell, a honeycomb pattern of α-tubulin was evident, not observed in wild-type, likely reflecting the cytoplasm remaining in the yolk cell that surrounds the yolk globules (Fig. 5C, S4B). In the wild-type sphere stage embryo, microtubules (MTs) and centrosomes were mainly organized around the nucleus or in the mitotic spindle apparatus of well-defined cells (Figs. 5D, S4B). In contrast, *kazu^p26thbd^* embryos displayed a severely altered blastoderm morphology at the same developmental stage, revealing a syncytial layer-like blastoderm expanded over the yolk cell, including diffuse MT distribution and a multitude of centrosomes (Figs. 5D, S4B).

These results indicate that blastomeres and blastoderm formation are affected in the early mutant embryo and suggests that Kazu is a critical factor in regulating the incorporation of cytoplasm into the developing blastodisc after egg activation. Thus, the *kazu^p26thbd^*mutant and other cytoplasmic segregation mutants (Dosch et al., 2004)(RF and MCM, unpublished) provide excellent genetic entry points to understand the molecular control of cytoplasmic volume acquisition and blastomere size regulation during early vertebrate embryogenesis.

### spotty regulates chromosome integrity, MTOC organization and microtubule nucleating activity during the oocyte-to-embryo transition

To determine the nature of the defects provoked by the *spotty* mutation, we investigated the development of the mutant egg and early embryo. Images of live one-cell stage *spotty* egg showed that at 40 mpf the blastodisc grew after the phase of massive transport of cytoplasm (Figs. 6A). Staining of the yolk cell with DiOC_6_, which labels the YGs due to their high lipid content, revealed multiple, small cytoplasmic domains in cortical regions of the yolk cell between peripheral YGs of the *spotty* mutant egg, unlike in wild-type (Fig. 6B). Together, these results point to a cytoplasmic subcellular organization defect in the *spotty* mutant zygote.

**Figure 6.**
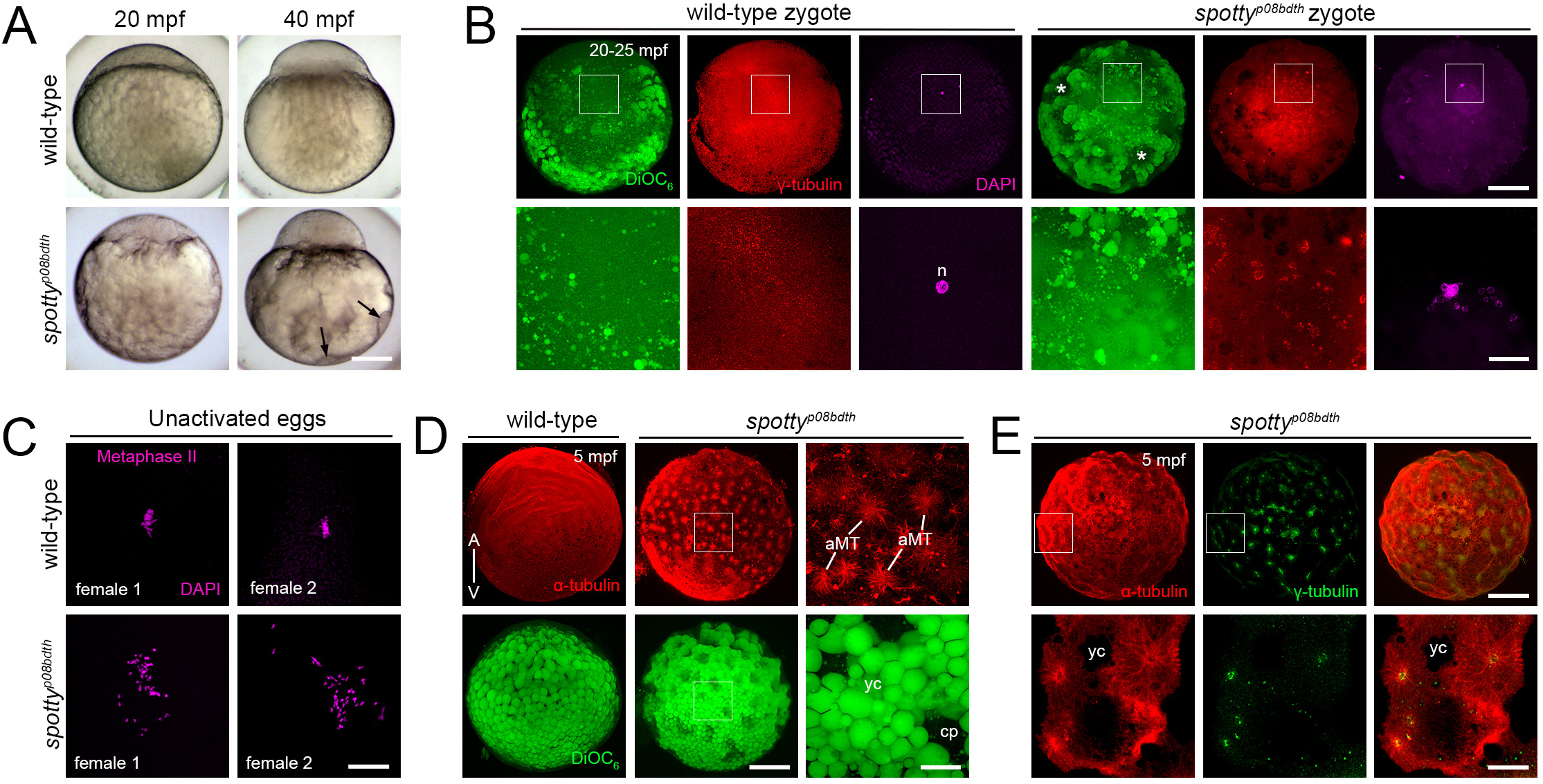
Chromosome, γ-Tubulin and microtubule organization in the *spotty^p08bdth^* mutant egg and zygote. **A.** Ectopic cytoplasmic domain formation in the *spotty* mutant. Instead of a regular organization of cytoplasm intermingled with YGs in the wild-type yolk cell (n=74 embryos from 2 females), the *spotty* mutant (n=65 embryos from 2 females) shows numerous, peripheral cytoplasmic domains (black arrows). **B.** Animal viewed confocal images of DiOC_6_– stained zygotes reveal that YG arrangement is affected, likely generating the peripheral patches of cytoplasm (white asterisks). Higher magnification DAPI– and γ-Tubulin-stained images show a perturbed nucleus formation and γ-Tubulin assembly in *spotty* (n=37 from 3 females) compared to wild-type (n=33 from 3 females) zygotes at 25 mpf. Boxes show magnified areas. **C.** DAPI-stained wild-type (n=20 from 2 females) and mutant (n=48 from 2 females) unactivated eggs showing chromosome organization during metaphase II of meiosis. **D.** α-Tubulin– and DiOC_6_-stained confocal z-projections showing MT and YG distribution at the cortex of wild-type (n=34) and *spotty* (n=41) eggs. Higher magnification images reveal the magnitude of such MT structures and YG organization in the mutant egg. **E.** α– and γ-Tubulin co-staining indicates that MTs emanate from numerous MTOCs across the mutant egg (n=14). Image 4 was selected from a z-stack containing 31 x 1 μm-thick optical sections across the mutant egg. Boxes show magnified (B, D and E) areas. bd, blastodisc; yc, yolk cell; yg, yolk globules; cp, cytoplasmic pocket; A, animal pole; V, vegetal pole; mpf, minutes post fertilization. Scale bar: 190 μm (A), 200 μm (B, top row), 50 µm (B, bottom row), 5 µm (C), 180 µm (D, low magnification), 38 µm (D, high magnification), 180 µm (E, top row), 60 µm (E, bottom row).

During the transition from meiosis to mitosis, the biogenesis of the centrosome and astral MT activity is coupled to zygotic nucleus formation and cell-cycle progression (Dekens, Pelegri, Maischein, & Nusslein-Volhard, 2003; Lindeman & Pelegri, 2012). The *spotty* phenotype shows abnormally cleaved cells (Fig. 1A), suggesting altered mitosis. Given that this process is accompanied by γ-tubulin assembly, and the nucleation and organization of MTs, we performed immunofluorescent staining of γ-tubulin and MTs immediately after fertilization. In the wild-type zygote, γ-tubulin is perinuclearly dispersed (Fig. 6B). However, the pattern of γ-tubulin distribution was dramatically altered in the mutant zygote with numerous aggregates localized around an unusual nuclear architecture in the blastodisc (Fig. 6B). Such nuclear organization suggested defective pronuclei behavior and nucleus formation. DAPI stained wild-type unactivated eggs revealed the arrested metaphase plate of meiosis II (Fig. 6C). However, several small DAPI spots, possibly dispersed individual or fragmented chromosomes were detected in *spotty* mutant unactivtaed eggs. This suggests that the maternal DNA integrity is likely compromised by a defective meiotic cycle in the oocyte. These results reveal that Spotty is essential to regulate: a) chromosome organization and segregation in the oocyte required for mononuclear assembly after fertilization, and b) the spatially restricted γ-tubulin localization during the first cell cycle.

In examining MTs in the egg, we found unusual MT arrays in the *spotty* mutant. Multiple unipolar aster-like MT structures were visualized at the cortex at 5 mpf. Higher magnification optical sections revealed apparent astral-like MT organization (Fig. 6D). At the same developmental period, the localization pattern of γ-tubulin confirmed that these MT-generating structures in the *spotty* mutant were indeed microtubule organizer centers (MTOCs) (Fig. 6E). The striking γ– and α-tubulin organization phenotypes in this mutant suggest that the factor encoded by the *spotty* gene suppresses MTOC organization, MT architecture and/or dynamics, in the egg.

We therefore examined the γ-tubulin localization in the *spotty* mutant during oogenesis. Immunostaining of isolated oocytes from adult ovaries, failed to detect any γ-tubulin foci in the wild-type diplotene stage I oocyte (>40 μm diameter) cytoplasm, consistent with expected centrosome elimination (Fig. 7A)(Elkouby, Jamieson-Lucy, & Mullins, 2016), a conserved feature of most animal oocytes examined. In contrast, several γ-tubulin spots were distributed throughout the cytoplasm of *spotty* mutant oocytes (Fig. 7A). As oogenesis proceeded, the number of γ-tubulin aggregates increased in the mutant oocyte (Fig. 7A,B). The ectopic supernumerary γ-tubulin aggregate phenotype was also found in cryosectioned adult ovaries (Fig. 7C).

**Figure 7.**
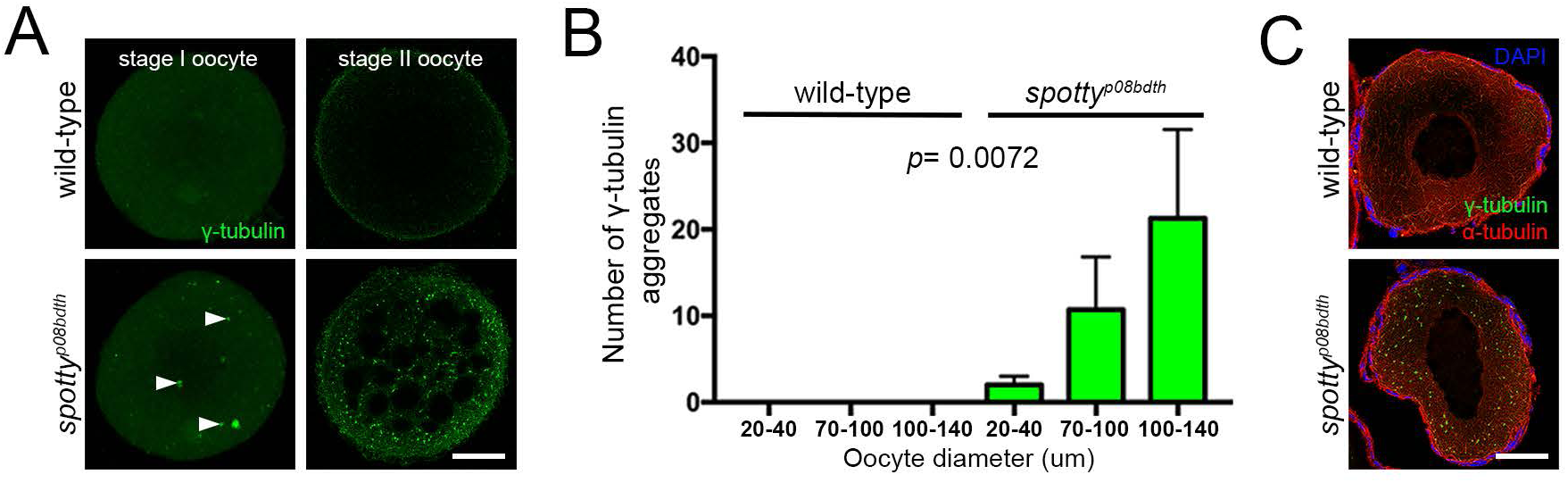
Ectopic supernumerary γ-tubulin aggregates phenotype in the mutant oocyte. **A.** Confocal z-projections (stage I) and optical section (stage II) of lateral viewed isolated wild-type (left) and *spotty* (right) oocytes showing the organization of cytoplasmic γ-tubulin aggregates (white arrow heads). **B.** Bars graph showing the γ-tubulin puncta number counted in three different size ranges of isolated oocytes: 20-40 μm (n=10 wild-type oocytes; n=10 mutant oocytes), 70-100 μm (n=11 wild-type oocytes; n=6 mutant oocytes), and 100-140 μm (n=10 wild-type oocytes; n= 9 mutant oocytes). **C.** Criosectioned whole ovaries showing the distribution of α– and γ-tubulin in a stage I wild-type (top, n=9) and *spotty* (bottom, n=8) oocyte. Scale bar = 25 μm (A, left column), 50 μm (A, right column) and 40 μm (C).

Altogether, these findings reveal severe defects in MTOC biology in the *spotty* mutant, thus representing the first described vertebrate mutant gene that regulates MTOC number. The identification of the *spotty* gene offers the opportunity to simultaneously understand the mechanism of how MTOC number is regulated in the oocyte and egg, and spatially organized through maternal control. And how MT dynamics are regulated by this factor during the oocyte-to-embryo transition.

## DISCUSSION

### Maternally controlled egg-to-embryo transition

The maternal-effect *krang*, *kazu*, *p28tabj,* and *spotty* mutants showed overtly normal development of the oocyte, including maturation of the cytoplasm and nucleus. However, these mutants did not successfully undergo processes triggered by egg activation. The distinct defects observed in these mutants are consistent with oocyte maturation regulated independently of aspects of egg activation. By mutant phenotyping, we have identified the essential role of the affected genes in the proper functioning of egg activation and early embryo formation. These findings provide valuable insights into the intricate mechanisms governing the egg-to-embryo transition in vertebrates. Understanding the precise molecular pathways and regulatory factors involved in this critical transition during embryogenesis has important implications for fertility and reproduction.

Our adult genetic screen not only contributes to the existing knowledge in the field but also sets the basis for future investigations. Thus, the identified maternal-effect mutant genes represent key players in coordinating and executing events in the egg to embryo transition. However, further studies are needed to fully elucidate the specific functions of these maternal genes and their participation in egg activation and subsequent embryonic development.

### Spatial and temporal control of maternal gene function in early embryogenesis

*kazu*, *p28tabj* and *spotty* mutant phenotypes suggest that while chorion elevation and blastodisc formation are both triggered by egg activation, they can be genetically separated and linked to the formation of compartmentalized cytoplasmic domains. This supports a model in which regionalized and distinct or divergent pathways in the egg regulate these aspects of egg activation. While separable genetically, other maternal-effect mutants show defects encompassing all or many aspects of egg activation, likely reflecting components regulating early steps of egg activation. For example, *brom bones/ptbp1a* is defective in inositol 1,4,5-trisphosphate (IP_3_) production and the Ca2+ wave that triggers egg activation, thus affecting all aspects of egg activation (Eno, Solanki, & Pelegri, 2016; Li-Villarreal et al., 2015; Mei et al., 2009). The maternal-zygotic *dachsous 1b* mutant exhibits slower activation of the egg, slower chorion elevation, as well as slower cell cleavage and early development (Eno et al., 2016; Li-Villarreal et al., 2015; Mei et al., 2009). The nature by which this atypical cadherin regulates egg activation processes and early embryonic development broadly is unclear. The maternal-effect *aura/mid1ip1l* mutant alters only a subset of egg activation processes with its primary function in regulating the cytoskeleton (Eno et al., 2016; Li-Villarreal et al., 2015; Mei et al., 2009). Hence, downstream of Ptbp1a and IP3, Dachsous and Aura, likely act in divergent egg activation pathways. Our findings of mutants affecting either CG biology or cytoplasmic segregation further show that these egg activation-associated events are also regulated independently.

### Krang: an uncharacterized, conserved factor regulating cortical granule biology

Following exocytosis, the contents of CGs modify the vitelline envelope causing it to expand, enlarging the perivitelline space, which is a well-characterized phenotypic trait to analyze high-quality eggs in ART (Balaban & Urman, 2006; Cappa et al., 2018; de Paola, Bello, & Michaut, 2015; Ten, Mendiola, Vioque, de Juan, & Bernabeu, 2007; Zhou et al., 2014). In addition, mutants with compromised CG exocytosis or vitelline envelope formation generate eggs with smaller perivitelline space compared to wild-type (Eno et al., 2016; Hau et al., 2020; Kanagaraj et al., 2014; Mei et al., 2009) or delay enlargement of the space (Li-Villarreal et al., 2015). Therefore, insight into the genetic program governing CG biology and vitelline envelope formation may provide biological markers for evaluating oocyte and egg quality for human ART. Recent findings provide new insight into the regulation of CG maturation and exocytosis during the vertebrate egg-to-embryo transition (Bello et al., 2016; Cappa et al., 2018; Cheeseman, Boulanger, Bond, & Schuh, 2016; de Paola et al., 2015; Kanagaraj et al., 2014; Mei et al., 2009), this work). However, a more comprehensive understanding of the molecular basis of how maternal factors regulate CG biogenesis, dynamics and content release remains to be revealed.

The loss of Krang function impairs egg activation, specifically affecting chorion elevation. We do not know the nature of Krang’s function in modulating secretory granule biology during female reproduction. Clues for its function are revealed by *krang* transcript localization, which is closely associated with CGs, strongly supporting a spatially-restricted function for this factor to CGs. These findings might shed light on maternal regulatory machinery that localizes translation, possibly to facilitate protein localization and function to, or within, CGs to regulate their content or function in CG exocytosis during egg activation. To understand Krang’s function in CG biology will require further investigation.

The human ortholog of Krang, KIAA0513 (71% identity), is predicted to function during neural development and its transcript to be dysregulated in individuals with schizophrenia and Alzheimer disease (Lauriat et al., 2006; Zhu, Jia, Li, & Jia, 2020). The *krang*/*kiaa0513* transcript is also zygotically expressed in the zebrafish hatching gland and nervous system (http://zfin.org). Indeed, the identification of *KIAA0513* splicing variants in the brain of related vertebrates, including the zebrafish, has also brought to light a possible exclusive and tissue-specific function of Krang in neural development (Lauriat et al., 2006; Zhu et al., 2020). If Krang acts in an oocyte-specific manner in vertebrates remains to be explored but in zebrafish its function is only essential maternally. In reported phenotypes from the International Mouse Phenotyping Consortium (IMPC), a knockout mutation of *Kiaa0513* (called 6430548M08Rik) yields homozygous mutant young adult males and females that had a low penetrance morphological defect of the kidney (1/16) and lymph node (2/16) (https://www.mousephenotype.org/data/genes/MGI:2443793). Thus, like in zebrafish, mouse *Kiaa0513* loss-of-function is homozygous adult viable and it is expressed in reproductive tissues in mouse and in the ovary in humans (https://www.genecards.org/cgi-bin/carddisp.pl?gene=KIAA0513#expression-protein), like *krang*. These results are consistent with a possible conserved role for Krang/KIAA0513 in CG biology in mammals, as in zebrafish in regulating vitelline envelope enlargement in egg activation during the egg-to-embryo transition.

### Maternal determinants of cytoplasmic organization during early embryogenesis

Following fertilization broadly in animals, early embryonic cells divide and maintain the total cytoplasmic volume inherited prior to the first cell cycle. These early cells are also unusually large and subcellular structures, such as the nucleus and mitotic spindle scale to a large extent to the larger cell size (Abrams et al., 2012; Good, Vahey, Skandarajah, Fletcher, & Heald, 2013; Hara & Merten, 2015; Huber & Gerace, 2007; Neumann & Nurse, 2007; Telley, Gaspar, Ephrussi, & Surrey, 2012). Despite these findings, how the total volume of cytoplasm is regulated and how a cell intrinsically distributes its cytoplasmic components remains poorly understood. We identified the *kazu^p26thbd^* mutant, which displays defects in blastodisc volume acquisition and small cells, thus representing a regulator of cytoplasmic domain formation during early embryogenesis.

After egg activation, a secondary and slower calcium transient observed in the central egg region coincides with and signals the spatial reorganization and animal-directed flow of cytoplasm (Fuentes & Fernandez, 2010; Sharma & Kinsey, 2008). It is possible that Kazu acts in blastodisc size acquisition by regulating the rate of the animal-ward cytoplasmic flow from the yolk cell, possibly in response to a calcium signal to appropriately allocate the subcellular yolk cell cytoplasmic domains to the zygote (reviewed in (Fuentes et al., 2018)). Actin, MTs, molecular motors and their regulators have been implicated in cytoplasmic segregation and RNA translocation through inhibitor and genetic studies (Fernandez, Valladares, Fuentes, & Ubilla, 2006; Fuentes & Fernandez, 2010; Gore & Sampath, 2002; Leung et al., 1998, 2000; Li-Villarreal et al., 2015; Shamipour et al., 2019). Therefore, it is possible that *kazu^p26thbd^*and other cytoplasmic segregation mutants disrupt upstream regulators or downstream targets of Actin/Myosin contractility and/or MT dynamics during egg activation. Ultimately, these cytoplasmic transportation defects may affect the arrival of maternal determinants to the developing blastodisc (Fuentes & Fernandez, 2010; Jesuthasan & Stahle, 1997; Tran et al., 2012). Future transcript localization analysis in *kazu^p26thbd^*, and other cytoplasmic segregation mutants, would test this hypothesis. Cell survival critically depends on cell volume regulatory mechanisms. Impaired steady-state cell volume regulation underlies physiological and pathological abnormalities such as cancer and hypertrophy (reviewed in Baltz & Tartia, 2010; Lloyd, 2013; Yang & Xu, 2011). In zebrafish, mutant genes that result in small blastodisc and blastomere phenotypes are candidates for contributing to the control of cytoplasmic domain formation and cell volume (reviewed in Fuentes et al., 2018). Thus, the *kazu^p26thbd^* mutant provides the opportunity to study the maternal control of embryo spatial distribution of cytoplasm and cell volume acquisition.

### Maternally-controlled MTOC assembly and microtubule nucleating activity during early embryogenesis

In most animal cells, the centrosome orchestrates the nucleation of both cytoplasmic MTs and the mitotic spindle during interphase and mitosis, respectively (Bornens, 2012; Conduit, Wainman, & Raff, 2015; Nigg & Stearns, 2011; Rieder, Faruki, & Khodjakov, 2001). During meiosis in most animals, however, an acentrosomal MTOC regulates the meiotic divisions, due to elimination of the maternal centrosome during oogenesis. Despite their importance as MTOCs, the genetic regulation of many aspects of MTOC and aMTOC assembly, structure, composition and function remain elusive. Intriguingly, multiple MTOCs/centrosomes are a hallmark in a growing list of human cancers, emphasizing the importance of regulating centrosome properties and number in health and disease (Chan, 2011; Gonczy, 2015; Harris, 2008). Remarkably, our understanding of how MTOCs acquire the ability to control their copy number and orchestrate the nucleation of cytoplasmic MTs, impacting on genome stability and embryogenesis progression, is poorly understood. During meiosis and early embryogenesis, little is known about the genetic program controlling how the oocyte, egg, and early embryo: a) transitions from centriolar to an acentriolar MTOC, and then back to a centriolar MTOC in the embryo, and b) keep MTOC *de novo* biogenesis repressed. This may be a particularly challenging task in the growing oocyte and egg, which is chock full of the factors needed to generate the multitude of centrosomes functioning during early cell divisions prior to zygotic genome activation. There are few known regulators of centrosome elimination and copy number control. In *Drosophila*, centrosome elimination is determined by Polo kinase activity and the reduction of pericentriolar material (PCM) components, which causes centriole loss before meiosis initiates (Pimenta-Marques et al., 2016). In echinoderms such as starfish, centriole reduction occurs during female meiosis, coupled to polar body formation, and under PCM loss regulation (Borrego-Pinto et al., 2016). This suggests that redundant mechanisms of centrosome elimination exist, and their genetic programs are executed temporally during meiosis in metazoan organisms. We do not know, however, if the ectopic MTOCs in *spotty* represent centrosomal or acentrosomal MTOCs.

The excessive number of aster-like MT structures and γ-Tubulin-positive aggregates in the *spotty* mutant oocyte and egg suggest that Spotty may function in suppressing centrosome and/or MTOC number, nucleating activity and ultimately, MT organization during the oocyte-to-embryo transition. Spotty may block protein recruitment and/or anchoring proteins essential for establishing MTOCs after oogenesis. Loss of centrosome-associated factors correlates with centrosome self-organization defects including centrosome amplification in cancer (Chan, 2011; Gonczy, 2015). Therefore, inhibiting this amplification effect is an attractive therapeutic target for tumor malignancy treatments. Determining the molecular identity of the *spotty* gene may identify a new repressor of supernumerary MTOCs, possibly centrosomes, and MT polymerization during the oocyte-to-embryo transition. However, the origin of the defect in the *spotty* mutant will require further investigation. Importantly, as the genetic network of *de novo* centrosome biogenesis is largely unknown, identifying novel molecular players will set the basis for future studies of the biological contribution of the centrosome/MTOC in human development and amplification in cell malignancy. Here, we present a genetic and functional model for the *in vivo* study of MTOC copy number regulation and aster-like MT control during early vertebrate embryogenesis.

## Materials and Methods

### Fish Mutagenesis and screening

Mutagenesis, mutant lines screening, and propagation approaches are described in the companion article (Fuentes et al., 2023).

### Mutations mapping and *krang^p30ahub^* gene lesion identification

The egg-to-embryo transition mutations were mapped as described in the accompanying paper (Fuentes et al., 2023). The *krang^p30ahub^*, *spotty^p08bdth^*, and *kazu^p26thbd^* mutations were mapped to chromosome (Chr) 18 (between 33 and 47 cM), Chr 2 (between 44 and 62 cM), and Chr 8 (between 95 and 104 cM), respectively (Table 1). For the *krang^p30ahub^* mutation, we generated the CR925798-1 marker for fine mapping and along with the Z58289 SSLP flanking marker (https://zfin.org/ZDB-SSLP-010801-361#summary), we narrowed the region to a ∼800 kb interval. Chromosomal locations of the SSLP markers were determined using the zebrafish reference genome Zv9.

For next generation sequencing, enriched and captured sequences (∼800 kb) were extracted from genomic DNA (gDNA) of wild-type and *krang^p30ahub^*females as previously described (Gupta et al., 2010). The libraries were then sequenced using an Illumina Genome Analyzer II.

### Generation of *krang^pΔ14^* mutant

Genome editing was carried out by targeting *krang*/*kiaa0513* in exon 5. A CRISPR target site 5′-gattgacagATGTCACCAGGTGG-3′ in the intron 4 and fifth coding exon joint of *krang*/*kiaa0513* was selected. The sgRNA construct was purchased from the University of Utah Mutation Generation and Detection Core. For injection, the synthetized sgRNA was incubated with Cas9 protein (PNA Bio, Cat# CP02-250) to form an RNP complex. 1-3 nl of 2.5 μM RNP complex were injected to 1-cell stage embryos. To evaluate cutting efficiency, we performed HRM (High Resolution Melt assay) analysis on F0 single injected embryos using MeltDoctor HRM Master Mix (Applied Bio-systems). Optimized flanking primers for HRM are indicated in Table S1. Remaining injected embryos were raised to adulthood to produce F1 families. To identify the *krang*/*kiaa0513* molecular lesion, we sequenced the amplified PCR product from transheterozygous fish genomic DNA using primers flanking the CRISPR/Cas9 target site (Exon 4-Intron 5, Table S1).

### Mutant lines genotyping and gene complementation test

*spotty^p08bdth^ and kazu^p26thbd^* alleles were genotyped using flanking SSLPs designed for positional cloning. *krang^p30ahub^*and *krang^pΔ14^* alleles were genotyped using <u>K</u>Biosciences Competitive <u>A</u>llele-<u>S</u>pecific <u>P</u>CR genotyping system (KASP, KBiosciences) (see Table S1).

For the complementation test, a *krang^Δ14^* heterozygous female was crossed to a *krang^p30ahub^* homozygous male to generate transheterozygous adult fish. Five transheterozygous and sibling *krang^p30ahub^* heterozygous females each were tested for phenotype. All transheterozygotes produced mutant embryos with small chorions, whereas sibling heterozygotes produced wild-type embryos. All transheterozygous females were tested at least twice and up to 5 times, reproducibly showing the mutant phenotype.

### *krang* transcript *in situ* hybridization and CG labeling

Fluorescent *in situ* hybridization of *krang* (this work) was performed in whole-mount oocytes and cryosectioned ovaries as described in Fuentes et al., 2013. For fluorescent mRNA detection, the Anti-POD antibody (1:500; Roche) and TSA system (PerkinElmer) were used. The *krang* digoxigenin-labeled probe was synthesized *in vitro* using DIG RNA labeling kit (Roche) with T3 (sense) or T7 (antisense) polymerase from zebrafish *kiaa0513* full length cDNA sequence cloned into the pBluescript II SK+ vector.

CG staining and number quantification were carried out in wild-type and *krang^p30ahub^* mutant unactivated and activated eggs as described in the accompanying paper (Fuentes et al., 2023).

### Whole-mount immuno– and actin staining, and histology

For microtubule/centrosome and actin labeling, egg, zygotes, and early embryos were fixed, washed and blocked as in (Gard, 1991) and (Becker & Hart, 1999), respectively. For staining of criosectioned tissue, ovaries were dissected, fixed, and stored in 30% sucrose as described (Escobar-Aguirre, Zhang, Jamieson-Lucy, & Mullins, 2017). Primary antibodies: mouse anti-α-tubulin (1:500, Sigma) and rabbit anti-γ-tubulin (1:2000, Sigma). Secondary antibodies: anti-mouse IgG1, or anti-rabbit IgG, Alexa 488, Alexa 594, Alexa 633 (All 1:1000, Molecular Probes). To label actin, TRITC-Phalloidin (Sigma) was diluted to 0.5 μg/ml in PBT. When needed, samples were counterstained with DiOC_6_ (5 µg/ml), then washed in 1X PBS 4 times for 10 min each and mounted in Vectashield mounting medium with or without DAPI on glass slides.

Whole ovaries from wild-type and *krang^p30ahub^* females were dissected for H & E stained histological sections as in the companion article (Fuentes et al., 2023). Microscopy, imaging and image processing were carried out as also described in this issue.

### Pronase treatment

Fertilized *krang^30ahub^* eggs from 2 females were exposed to 1.5 mg/ml pronase solution in 1X E3 buffer for 10 minutes at 28°C. Control eggs were incubated in 1X E3 pronase-free buffer. To finish dechorionation, embryos were washed with 1X E3 buffer three times by gentle trituration using a Pasteur pipette, and examined at gastrula stage under a stereomicroscope. Cleaving rate was calculated using the Prism 7 (GraphPad) software.

### Electron microscopy

For transmission electron microscopy, whole wild-type and *krang^p30ahub^* ovaries were dissected and fixed in 2.5% glutaraldehyde, 2.0% paraformaldehyde, and 0.1M sodium cacodylate overnight at 4 °C. Washes, post-fixation, dehydration, embedding, thin sections, and staining were performed by using standard methods at the UPenn EM core facility. Stained sections were examined with a JEOL JEM 1010 electron microscope.

### γ-tubulin puncta number analysis

Maximum intensity projection images of different sized isolated oocytes were used to count cytoplasmic aggregates (γ-tubulin-fluorescent puncta) using Fiji software. Three different size ranges of oocytes were analyzed: 20-40 μm, 70-100 μm and 100-140 μm. Sample size is mentioned within the figure legend. Statistical analysis was performed using Prism 7 (GraphPad) software. Data are represented as mean ± SEM and analyzed with the one-way ANOVA test.

### Homology modeling of the zebrafish maternal Krang/Kiaa0513 factor

The structure prediction of the zebrafish Krang/Kiaa0513 protein, is described in the companion article (Fuentes et al., 2023). Neighbor joining protein sequence-based phylogenetic tree of Krang/Kiaa0513 homologs was handled using FigTree v1.4.2 (http://tree.bio.ed.ac.uk/software/figtree/). The tree was reduced from its original size by allowing a maximal sequence difference (Max Seq Difference) in fraction of mismatched residues of 0.85, resulting in 179 sequences. Bootstrap values from 500 iterations are shown for each node in the resulting cladogram (Fig. S2).

## Acknowledgments

We thank Drs. Leonardo E. Valdivia, Matías Escobar-Aguirre and Matthew Good for valuable comments on the manuscript; Dr. Juliane Bremer for kindly providing the pBluescript II SK+ vector; Dr. Andrea Stout and the CDB (University of Pennsylvania), and CMA Bio-Bio (Universidad de Concepcion) microscopy cores for assistance in the use of the confocal microscopes; Amy Kugath and fish facility staff (University of Pennsylvania), and Andrea Aguilar and Karina Vega-Drake (Universidad de Concepción) for technical assistance and fish care; and the Next Generation Sequencing core at the University of Pennsylvania Perelman School of Medicine.

This work was supported by National Institutes of Health (NIH): R35GM131908, R21HD094096, and R01HD069321 to M.C.M. Becas Chile/Conicyt Proyecto Postdoctorado 74130048, Apoyo FCBI2019-01, Proyecto VRID Investigación Multidisciplinaria 220.031.117-M, and ANID Proyecto Fondecyt de Iniciación 11201118 to R.F. Damon Runyon Postdoctoral Fellowship DRG1826-04 to F.L.M.. NIH training grant T32-HD007516 and NRSA postdoctoral fellowships 5F32GM77835 to E.W.A, 1F32GM080926, and American Cancer Society postdoctoral fellowships PF-09-125-01-DDC to L.K, and PF-05-041-01-DDC to T.G.

