## Supplemental Table and legends for "Maternal regulation of the vertebrate egg-to-embryo transition"

**Table S1. Primers for *krang* mutant genotyping and gene mutations sequencing.**

| **Primer** | **Sequence** |
| --- | --- |
| CR925798-1 | Forward Primer: 5’-TTTTTCATTTACTGGTATTGAAAGTG-3’  Reverse Primer: 5’-TGTACAGAAATTGGGGGAAAA-3’ |
| *krang^p30ahub^* | 5’-GAGGAGAAGGCTCGCTTTGGAGAGCTCTGTACTGGA  GACAATGGGAAAGG[C/T]AGGGAATGGTTCTCCAAATAC  GTCAGTGCGCAACGCTGCCAC-3’ |
| HRM | Forward Primer: 5’-TGGTGCAGTCTTTCGCTGTT-3’  Reverse Primer: 5’-AGGTGTTTGGCAGGGCAATA-3’ |
| *krang*_Exon4-Intron5-6 | Forward Primer: 5’-CGCTGCCACTCCAAATG-3’  Reverse Primer: 5’-GACACAATAATGGGCAGTACC-3’ |

**Supplementary Figure Legends**

**Figure S1. CGs and embryonic phenotype in *krang* mutants, chorion structure analysis, and the mutation consequence. A.** Confocal z-projections (60 μm depth) of acid fixed and MPA stained wild-type and mutant activated eggs. MPA staining reveals intact CGs of wild-type and mutant eggs at 20 mpa. CGs persist in *krang* eggs, revealing that the release of their content is compromised. In addition, numerous small CGs were retained after activation in eggs from these mutant females. **B.** Curves showing CG exocytosis rate in wild-type and *krang* mutant eggs. Data are means ± SD. Time constant (τ) for wild-type and mutant was 11.67 and 13.23, respectively. **C.** Ultrastructural analysis of the chorion morphology in wild-type (n=12) and *krang* (n=14) oocytes. Boxes show magnified areas revealing different electron density aspects of the chorionic zones I, II and III. **D.** *krang* mutation results in a frameshift in the coding sequence, which generates a premature stop codon (in red) following 24 aberrant amino acids and a truncated protein. Scale bar= 40 µm (A), 8.6 µm (C, left column), 2 µm (C, right column).

**Figure S2. Evolutionary fate of Krang across the animal kingdom.** Cladogram generated by analyzing Krang amino acid sequence showing its phylogenetic conservation among metazoans. Color coded organisms are representative members of invertebrate and vertebrate phyla/classes. The numbers at the bases of the branches indicate bootstrap values obtained from 500 iterations. The scale bar on the bottom represents distances in residue substitutions per site.

**Figure S3. CRISPR/Cas9-based mutation of zebrafish *krang* perturbs egg activation and embryo development. A.** Schematic of the zebrafish *krang*/*kiaa0513* locus and CRISPR/Cas9 targeted region (red rectangle). Exons are shown as green boxes and introns as black lines. Sizes are not to scale. The 14 nt deletion and premature STOP codon generated by the mutation are shown. **B.** Representative images of mutant early embryos laid by mutant females. A normal and defective chorion elevation in wild-type and mutant embryos, respectively, are indicated (black arrows). Notice that mutant early embryos display altered blastoderm formation. Scale bar= 311 µm (A).

**Figure S4. Characterization of the 1-day post-fertilization and cell morphology *kazu* phenotype. A.** *kazu^p26thbd^* early embryo exhibits a smaller blastodisc phenotype and a severely affected cytoplasmic segregation. Most *kazu^p26thbd^* mutant embryos die about 5 hpf and survivors may give rise to embryos with a reduced body. **B.** DAPI, α-tubulin, and centrosome staining showing abnormal blastoderm formation. Cell boundaries and mitotic figures are not observed in most of the mutant (n=25/33) compared to wild-type (n=30/35) embryos examined. Notice the formation of a syncytial nuclei layer in the 5 hpf *kazu^p26thbd^* embryo. Scale bar= 750 µm (A), 180 µm (B, left column), 150 µm (B, right column).
