## Supplementary figures and images for "Maternal regulation of the vertebrate egg-to-embryo transition"

### Supplemental Figures

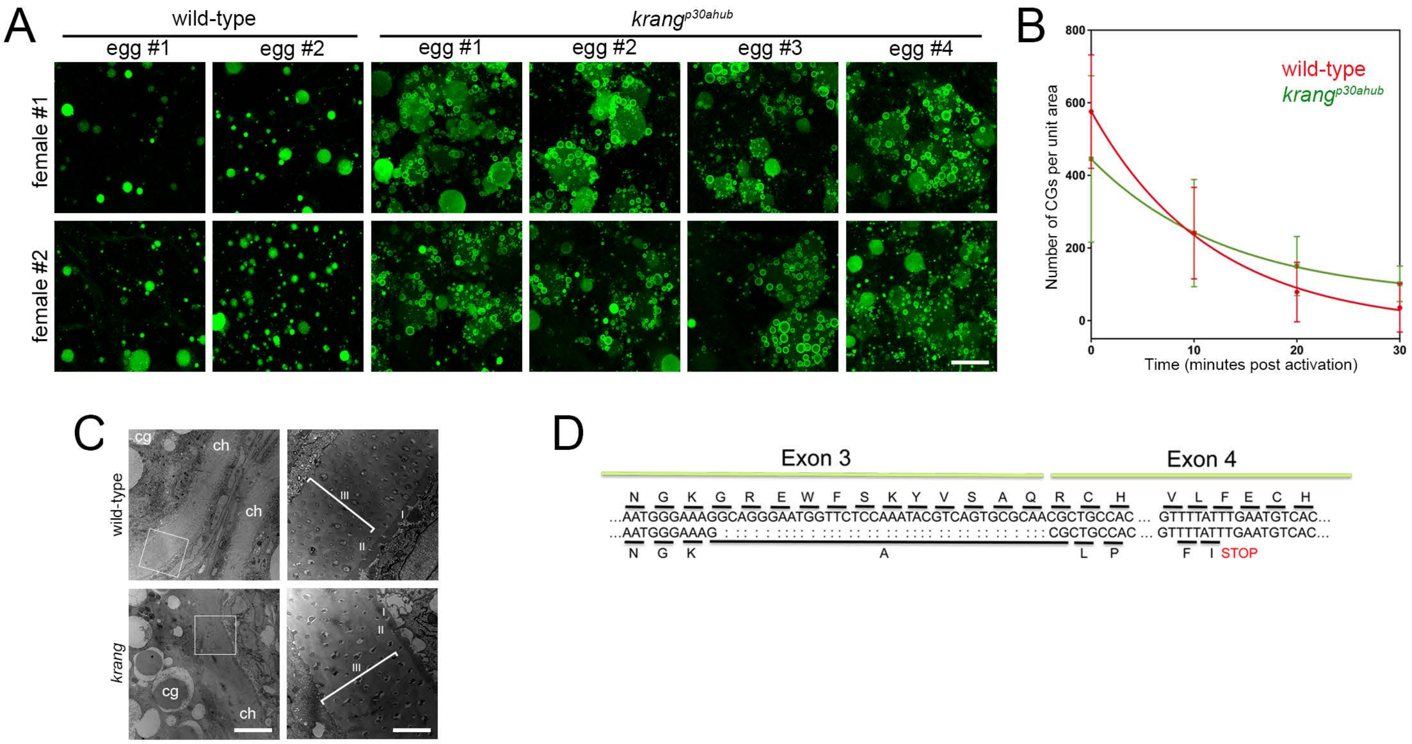

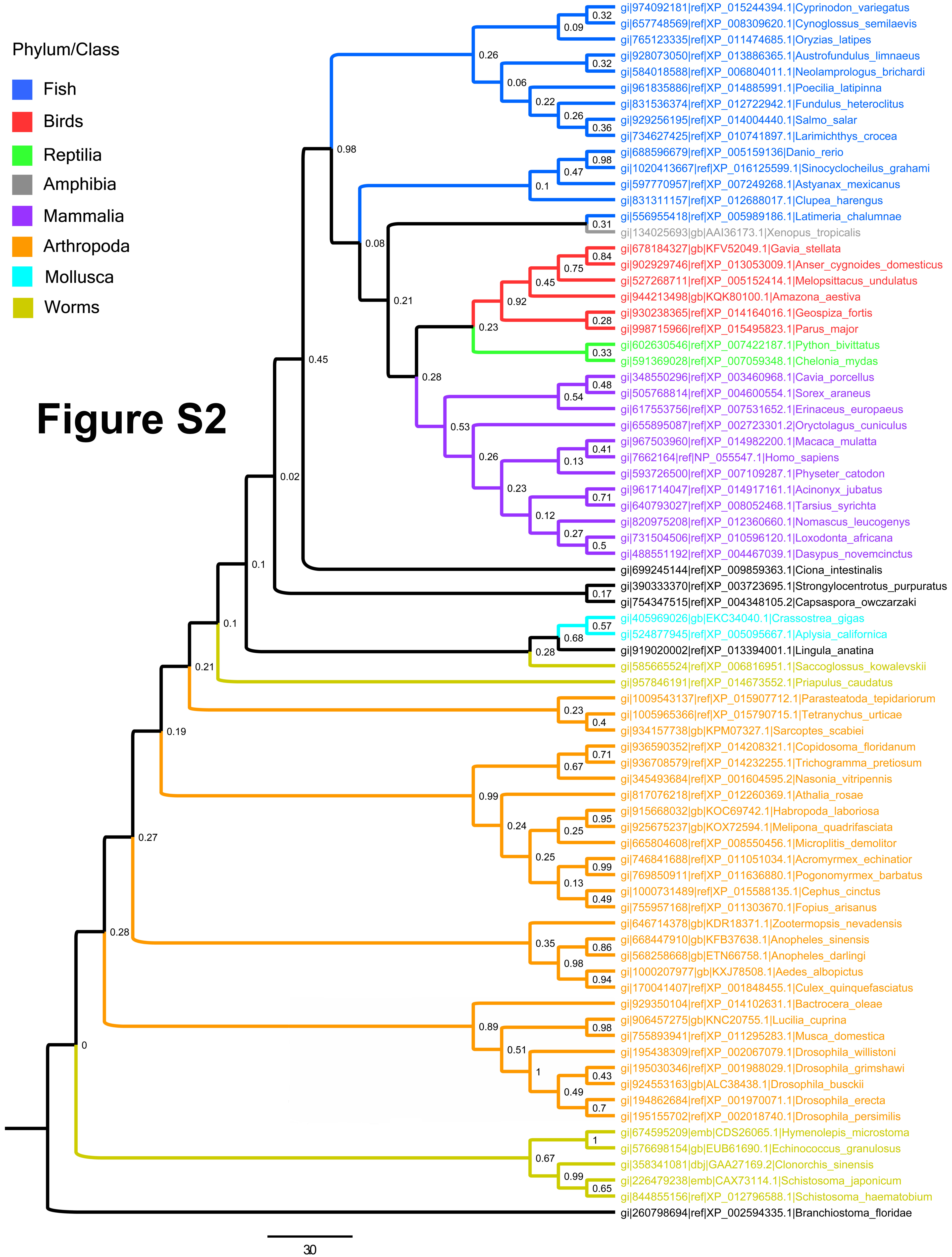

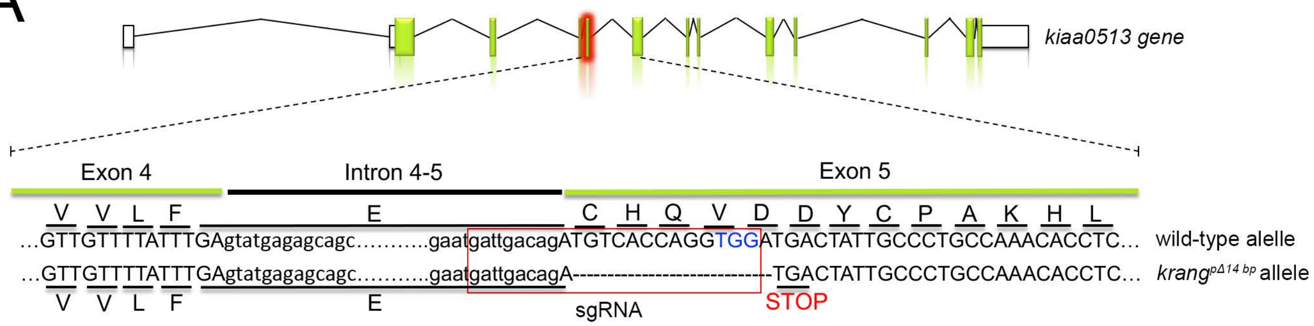

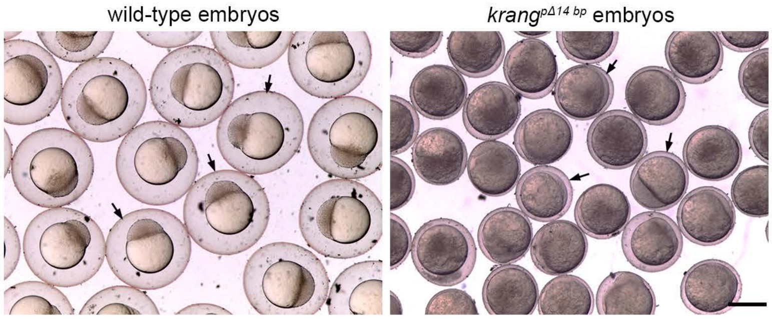

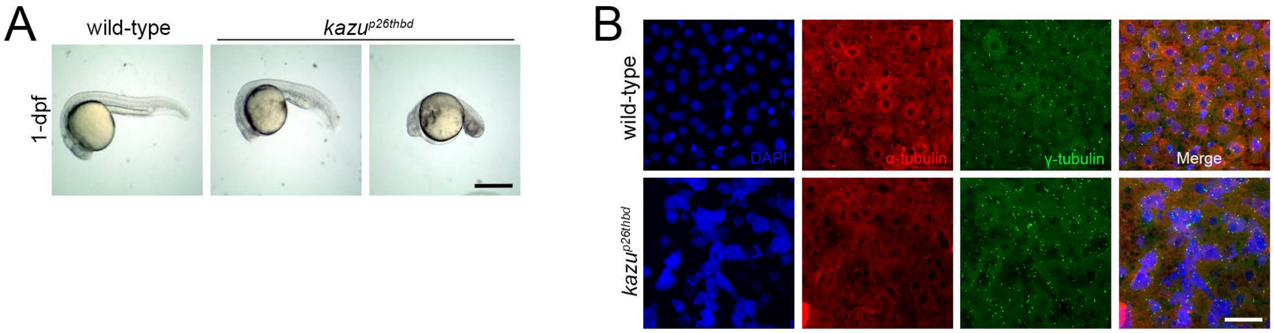
